# Neural Context Reinstatement of Recurring Events

**DOI:** 10.1101/2024.11.07.622553

**Authors:** Adam W. Broitman, Michael J. Kahana

**Author notes:** Correspondence concerning this article should be addressed to Michael J. Kahana,. Author Note The authors gratefully acknowledge support from National Institutes of Health grant MH55687. We thank David Halpern and Ryan Colyer for assistance with designing and programming the experiment, and Max Weinstein, Lucas Flahault, David Halpern, and Nathaniel Greene for assistance with data collection. The ideas and data appearing in this manuscript have not been disseminated before.

## Abstract

Episodic recollection involves retrieving context information bound to specific events. However, autobiographical memory largely comprises recurrent, similar experiences that become integrated into joint representations. In the current study, we extracted a neural signature of temporal context from scalp electroencephalography (EEG) to investigate whether recalling a recurring event accompanies the reinstatement of one or multiple instances of its occurrence. We asked 52 young adults to study and recall lists of words that included both once-presented and repeated items. Participants recalled repeated items in association with neighboring list items from each occurrence, but with stronger clustering around the repetition’s initial occurrence. Furthermore, multivariate spectral EEG analyses revealed that neural activity from just prior to the recall of these words resembled patterns of activity observed near the item’s first occurrence, but not its second. Together, these results suggest that the initial occurrence of an event carries stronger temporal context associations than later repetitions.

**Research Transparency Statement:** The authors report no conflicts of interest with respect to the authorship or publication of this article. This research was conducted with support from National Institutes of Health grant MH55687. The current study was not preregistered. Data are available upon reasonable request and with proper approval from the University of Pennsylvania research and ethics entities. Requests should be directed to Adam Broitman. Key EEG and behavioral data analysis scripts are available for download at https://github.com/awb99cu/repFRcode.Studymaterialsareavailablefordownloadathttps://memory.psych.upenn.edu/PEERS.

## Introduction

According to retrieved-context models of episodic memory, episodic recollection involves reinstating contextual information bound to specific prior events, leading to associations between events that happened close together in time (Howard and Kahana, 2002; Polyn et al., 2009). A rich literature has characterized the cognitive and neural processes involved in the encoding and retrieval of these associations (Manning et al., 2011; Polyn and Cutler, 2017; Smith et al., 2022; Folkerts et al., 2018; Lohnas et al., 2023). However, in everyday life, we frequently experience routine, recurring events that are nearly indistinguishable from one another (e.g., brushing one’s teeth), and most autobiographical memories are amalgams of multiple distinct experiences blended into what Ulric Neisser termed “repisodic” memories (Neisser, 1981). Although much work has explored the effects of repetition on memory, it remains unclear how context information from multiple events becomes associated with a single item. For instance, do people tend to reinstate the context of the initial appearance of an item, the most recent appearance, or a composite of all appearances? The current study aims to address this question by investigating the behavioral and neural markers of context reinstatement for repeated experiences.

Early cognitive theorists posited that repeating an item strengthens its memory trace, increasing its chances of later retrieval (Thorndike, 1934; Hull, 1935). Subsequent work demonstrated that individuals can separately recall details of each occurrence of a repeated event, suggesting that a single item can become bound to multiple context representations (Yntema and Trask, 1963; Hintzman and Block, 1971). Recent computational frameworks, such as the *Context Maintenance and Retrieval* model (CMR; Polyn et al., 2009; Lohnas et al., 2015), have further refined our understanding of repetition effects. CMR proposes that items encoded in a list become associated with a slowly drifting *temporal context* signal, which captures information about the sequences of events and elapsed time between them. During recall, reinstating an item’s bound context representation cues the retrieval of other items encountered at neighboring time points. CMR similarly predicts that repeating a word during encoding reactivates the temporal context of its initial presentation. As a result, the word becomes linked to a representation that merges the context of its initial presentation with the gradually evolving context of its later repetition. Thus, retrieving a repeated item should involve reinstating the context of both presentations.

Evidence supporting CMR’s predictions include findings that repetition effects strengthen as spacings between same-item presentations increase (Madigan, 1969; Melton, 1970). This suggests that increasing the distinctiveness of each of the item’s contextual cues improves its chances for retrieval. Later studies found that people are more likely to successively recall list items that immediately followed both presentations of a shared item (Howard et al., 2007; Lohnas and Kahana, 2014), indicating that even when items are presented at distant serial positions, both are still associated with the same repeated item. Other work has demonstrated that item repetition can cue “pattern completion”, or the retrieval of associative information that forms coherent event representations around that item (McClelland et al., 1995; Wills et al., 2005; Horner and Burgess, 2014). Aligning with these results, neural recordings have shown that repeating an item within a sequence evokes neural patterns similar to those observed at its initial occurrence, and that this effect is correlated with later memory for that item (Xue et al., 2010; Halpern et al., 2023). Although the bulk of this research suggests that repeated items are bound to multiple context representations, it does not indicate whether individuals reinstate one or multiple of these representations while recalling the items.

Various methods exist for measuring temporal context reinstatement during recall. Analysis of free recall responses can quantify *temporal contiguity*, or the tendency to successively retrieve items that appeared at neighboring serial positions, as an indicator of how strongly items become bound to their temporal contexts (Kahana, 1996; Polyn et al., 2009). More recent work has applied Represenational Similarity Analysis (RSA) to identify neural markers of context reinstatement. These studies reported that when an individual recalls an item, their neural activity is not only similar to that observed during the study of the item, but also resembles the activity from neighboring list items, with similarity decreasing as the serial position distance increases (Manning et al., 2011; Howard et al., 2012; Ten Oever et al., 2021; Lohnas et al., 2023). However, these metrics have thus far only been applied to measure context reinstatement for once-presented items.

The current study investigates temporal context reinstatement for repeatedly presented items in a multi-session verbal free recall experiment. Participants encoded lists of words presented either once within the list (Single items) or twice (Repeat items), and then recalled as many words as they could following a brief delay. We aimed to address two key questions regarding the recall of Repeat items: 1) Do individuals cluster their retrievals with items presented near the first or second occurrence of a repeated item? and 2) Can we detect neural evidence of temporal context reinstatement for the first, second, or both presentations of the item? We approached these questions by analyzing both behavioral and neural patterns of context reinstatement.

CMR assumes that repeating an item within a list will activate the temporal context of its prior presentation, thereby strengthening its associations with other items presented near its initial occurrence. As a result, participants would be more likely to cluster the retrieval of repeated items with other items near its first presentation and to reinstate neural activity from that initial occurrence. Supporting this hypothesis are classic A–B, A–C paired associate learning studies, which found that although A–C pairings are strongly retrieved immediately after they are presented, memory for A–B associations exceeds that of the later-learned A–C pairings following a 24 hour study-test delay (Briggs, 1954; Randel and Havrda, 1968; Underwood, 1957). This result suggests that repeating the presentation of an item (A) strengthens its associations with its previously learned paired associate (B). However, an alternative possibility is that recency effects will influence participants to reinstate items near the second presentation of Repeat words, which occur at later serial positions. People are more likely to remember repeated events that recurred recently (Means and Loftus, 1991), and this could lead individuals to preferentially reinstate the most recent occurrence. The current experiment will test these competing predictions by separately assessing temporal context reinstatement for each occurrence of a repeated item.

## Method

### Participants

We selected an *a priori* target sample size of 50 participants based on data from Halpern et al. (2023), which investigated study-phase retrieval of repeated items using intracranial EEG. Using G*Power 4.1 (Faul et al., 2009), we assumed an effect of size Cohen’s f = .4 with a power (1 – *β*) of .95 in an F-test of neural similarity between two same vs. different items presented during encoding. We subsequently recruited 52 native English speaking, neurocognitively healthy adults aged 18-30 (33 female, mean age = 22.9) from the University of Pennsylvania campus and the surrounding Philadelphia community using fliers and online promotions. Participants were paid a base rate of $12 per session, and were offered an additional bonus of up to $18 per session for minimizing their eye blinks and head movements during the session. We awarded a final $50 bonus to participants who completed all six sessions of the experiment. The University of Pennsylvania IRB reviewed and approved all recruitment and testing procedures.

### Experimental Materials and Equipment

Participants sat unconstrained in a normally lit Faraday enclosure, approximately 90 cm away from a ViewSonic 17” LCD monitor (1024×768 pixels, 75Hz refresh rate). We programmed the experiment using custom Python and Unity scripts.

We constructed word lists using a pool of 1638 nouns taken from the University of South Florida free-association norms (Nelson et al., 2004). Each list consisted of 12 words, half of which were presented one time (Single items), and half were presented twice, separated by at least three intervening word presentations (Repeat items). This resulted in a total of 18 word presentations per list. The first presentation of a Repeat item would always occur within the first 12 serial positions of a list, while the second presentation always occurred after the sixth serial position. Single items were presented across all serial positions. Using the Word Association Spaces (Steyvers et al., 2004), we constructed word lists such that adjacent words would have low semantic similarity to one another in order to minimize potential confounds between temporal and semantic associations among study items.

### Free Recall Experiment

In each of six sessions, participants completed 21 blocks of the verbal free recall task (see Figure 1). At the start of each block, the screen presented a fixation for seven-seconds, followed by a three-second countdown to the beginning of word presentation. Each word encoding trial (duration = 4100-4600 ms) consisted of a word presented at the center of the screen for 1600 ms. Following word presentation, the screen remained blank for a jittered (i.e., variable) interstimulus period of 2500-3000 ms. Following the presentation of the final list item, a 15 s post-encoding delay occurred, followed by an auditory tone and the appearance of a row of asterisks onscreen. At this time, participants had 30 s to vocally recall as many items as they could remember from the previous word list.

**Figure 1.**
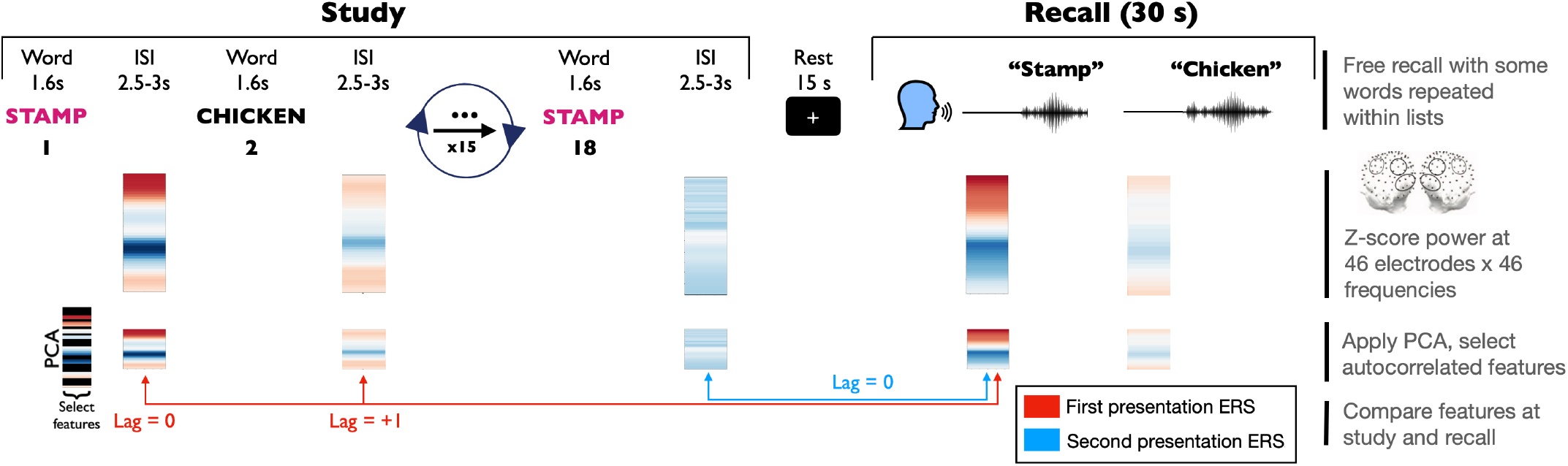
Overview of experiment and neural analyses. Participants encoded lists of 12 words presented for 1600 ms each, followed by a 2.5-3 s ISI. Half of the words (Single items) were presented once, and the other half (Repeat items) were presented twice throughout the list. Following a 15 s rest period, participants vocally recalled as many words as they could remember for 30 s. Following data collection, we computed spectral power at 46 frequencies with 46 electrodes using data from the ISI at each encoding event (1600-4100 ms), and from just prior to each retrieval (-900 to -200 ms). We performed PCA on the spectral wavelets from each encoding epoch, selecting components that explained a significant proportion of the variance in activity and that changed slowly across trials. Finally, we obtained encoding-retrieval representational similarity (ERS) values by computing the Pearson correlation coefficient between the encoding and retrieval epochs.

### EEG methods

#### Data collection and preprocessing

We collected EEG data using a 128-channel BioSemi ActiveTwo geodesic system at a sampling rate of 500 Hz. We instructed participants to minimize blinks and body movements during the experiment. Following data collection, we applied a Butterworth notch filter at 60 Hz and 120 Hz to eliminate electrical noise, as well as a low-pass filter at 1 Hz to eliminate slow drift artifacts. We identified bad channels as those with extremely high or low (|*z*| > 3) log-variance, or which had an extremely high (*z* > 3) Hurst exponent relative to other channels. After dropping the bad channels, we re-referenced the recordings to the common average across all remaining electrodes. To remove eyeblinks and other artifacts, we used an automatic artifact detection algorithm based on independent components analysis (Nolan et al., 2010).

#### Neural encoding-retrieval representational similarity (ERS) analysis

For each encoding and recall trial, we computed power spectral density at 46 logarithmically spaced frequencies ranging from 2-100 Hz using a Morlet wavelet transformation with a wave number of 5 and a 600 ms mirrored buffer. We selected a region of interest (ROI) consisting of 46 channels, in accordance with prior work that used similar EEG caps to detect multivariate spectral biomarkers of memory (Broitman et al., 2024; Lohnas et al., 2023; Li et al., 2024). Figure 1 topographically plots the ROI electrodes, which covers mid-frontal, parietal, and occipital scalp sites. We defined the encoding period as the ISI period from each trial (1600-4100 relative to stimulus onset). We selected this time window for two reasons: to account for any delay between the word’s appearance and the participant’s processing of the word, and to minimize any overlap in processing of the previously presented word. Neural ERS studies commonly use data from the ISI period for encoding-retrieval comparisons (Sederberg et al., 2007; Ten Oever et al., 2021; Yaffe et al., 2014; Howard et al., 2012), and others have implemented a delay following stimulus onset when selecting the encoding epoch (Manning et al., 2011; Lohnas et al., 2023; Liang and Preston, 2017; Gershman et al., 2013). For retrieval trials, we examined the time window from -900 to -200 ms preceding the onset of vocalization of the word. This epoch was chosen to minimize the potential for overlap of output of the previously recalled item and to omit motor artifacts from recalling the current item.

Among recall trials, we excluded those occurring within 1000 ms after a recall or vocalization to avoid overlap in activity between retrievals. Furthermore, mirroring the method of Lohnas et al. (2023), we excluded recalls from output position 1, as such recalls can reflect retrieval from immediate short-term memory rather than temporal context reinstatement (Kahana, 1996). If, during encoding, two repeat items occurred sequentially during both presentations, then these trials would be excluded from analysis to address the potential confound between sequence reinstatement and study-phase retrieval.

We followed methods used in prior studies to identify neural features that were autocorrelated, i.e., changing slowly across the encoding period (Manning et al., 2011; Lohnas et al., 2023). We first applied principle components analysis (PCA) to the log-transformed and Z-scored wavelet values. After excluding components with an explained variance regression score below 1.0, for each of the remaining components we computed the Pearson *r* correlation coefficient between adjacent encoding trials within a list. We then obtained a summary statistic for the component’s correlation across lists by taking the Fisher *z*-prime transformation *F* of each *r* value, and then combining them into a summary statistic 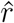 using the following formula:

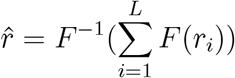

We also computed a summary measure 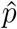 for *p*-values obtained from each Pearson correlation by summing the inverse normal-transformed *p*-values on the cumulative density function. We defined autocorrelated feature components as those with 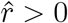 and 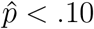. We excluded sessions that did not yield at least five feature components meeting our criteria for autocorrelation and explained variance. This process excluded 15 of the original 253 experimental sessions for a total of 238. All 52 original participants remained included in the final analysis.

We defined neural similarity between two events as the Pearson correlation coefficient between the corresponding neural features from each event. For each retrieval event, we computed its neural similarity with all encoding events from the same session. Because there were a total of 378 word presentations during each session (18 words *×* 21 lists), this resulted in an RSA matrix of 378 × *n*-recalls.

## Results

### Behavioral results

#### Recall rates of Single and Repeat items

Figure 2A presents the recall rates for Single and Repeat items. As predicted, participants recalled Repeat words more frequently than Single words, *t*(51) = 9.7, *p* < .001, replicating prior findings that recall rates increase with additional presentations (Tulving, 1967; Howe, 1970; Daves and Rinn, 1971).

**Figure 2.**
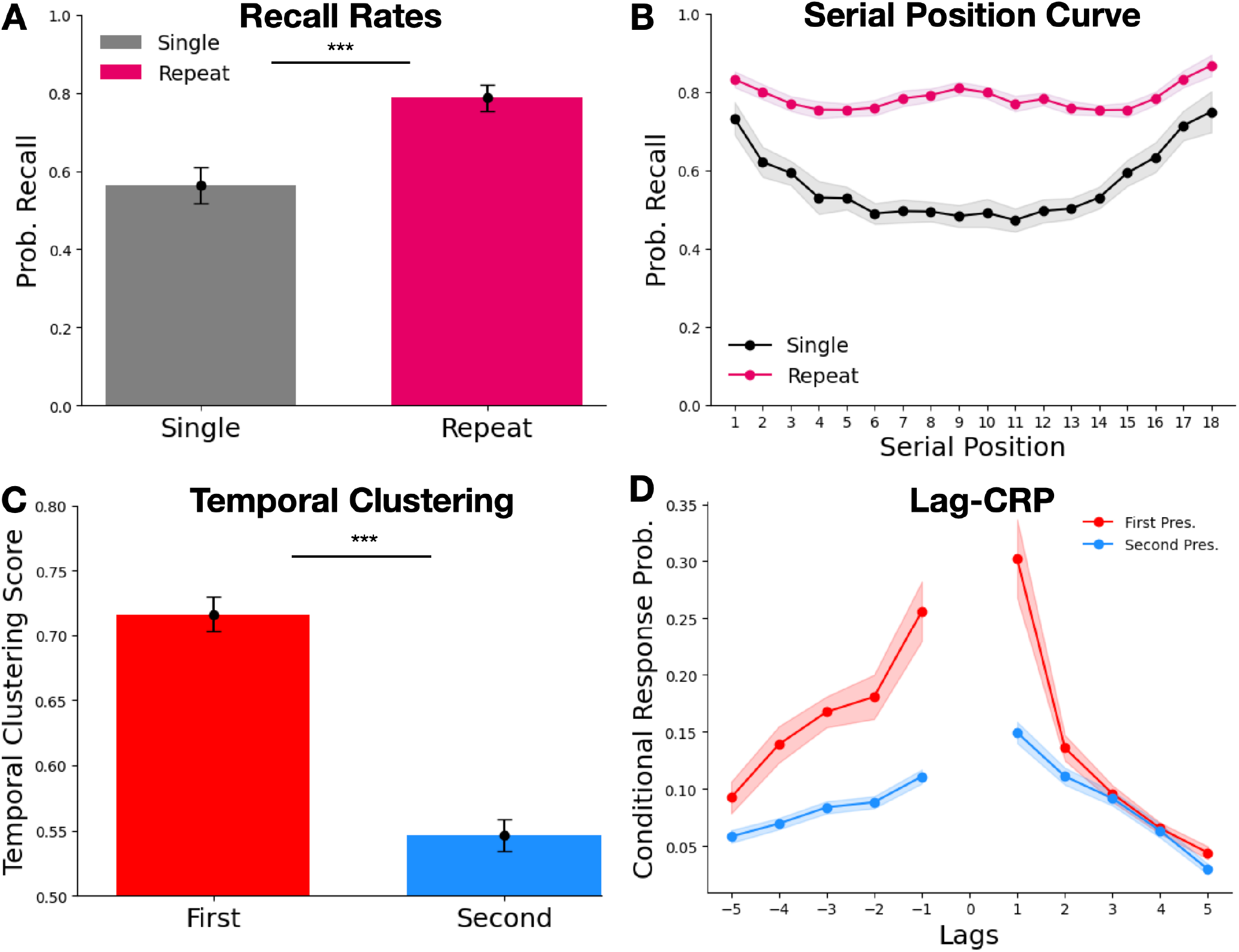
Behavioral Results. **A.** Recall rates for Single (gray) and Repeat items (purple). **B**. Serial position curve (SPC) for Single and Repeat items. Repeat item SPCs were computed by counting each presentation and recall toward both of the item’s serial positions. **C**. Temporal clustering scores for recall transitions from Repeat items attributed to the serial position of the first presentation (red) or second presentation (blue). **D**. Lag-conditional response probability function for recall transitions from Repeat items. Bar plot error bars indicate 95% standard error confidence intervals, and line graph error bands indicate 95% repeated-measures confidence intervals (Loftus and Masson, 1994).

The bottom panel of Figure 2B presents recall probability as a function of serial position (*serial position curve*; SPC) for both Single and Repeat items. To compute the SPC, we counted each repeat item for both of its corresponding serial positions. The SPC for Single items demonstrates standard primacy and recency effects, with elevated recall rates at early (SPs 1-3) and late (SPs 16-18) list positions, *t*(51)′*s* > 9.2, *p*′*s* < .0001, relative to middle serial positions 4-15. However, in addition to primacy and recency effects, the Repeat item SPC reveals elevated recall rates in the *middle* list positions (8-10), *t*(51) = 4.8, *p* < .0001, relative to SPs 4-7 and 11-15. We hypothesized that this could have arisen from primacy and recency effects. If items initially presented in these middle positions were usually repeated at the end of the list, then they might benefit from recency effects. Similarly, items initially presented during primacy positions might be likely to recur during these middle positions. To verify this, we examined all repeat items initially presented at positions 8-10, and found that the average serial position for the second presentation of these items was 16.7 (SD = 1.2). We similarly examined items whose second presentation occurred at SPs 8-10, finding that their first presentation occurred at an average SP of 1.9 (SD = 1.0). The elevated recall rate of Repeat items from middle serial positions therefore likely reflects primacy and recency effects originating from other presentations of those same words.

#### Behavioral evidence for context reinstatement of Repeat items

Our primary question was whether participants reinstated temporal context from one or both presentations while retrieving a repeated item. We therefore examined participants’ tendencies to cluster recalls of Repeat items with other list items presented near their first or second presentation. We computed the *lag conditional response probability* function (lag-CRP; Kahana, 1996), which plots the probability of successively recalling items as a function of their distance from one another in the study list. This distance is measured in *lags*, or the difference in serial positions between each successively recalled item. Polyn et al. (2009) updated this metric with the *temporal clustering score*. For each recall transition made, the temporal clustering score computes the proportion of lags to available items (i.e., any not-yet-recalled items from the study list) that the actual lag is *less than*. When averaged over all recall transitions, this results in a score ranging from 0 to 1, with larger numbers indicating a greater degree of temporal clustering. For each Repeat item *i* that was recalled, we separately computed the transition lag for both of its serial positions *i*_1_ and *i*_2_. Following the method first presented by Kahana and Howard (2005), if the word recalled after item *i* was also a Repeat, we would select whichever serial position minimized the absolute lag to the serial position being transitioned from. This procedure produced separate lag-CRP functions and temporal clustering scores for the first and second presentations of Repeat items (Fig. 2C and D). If no temporal ordering effects were present, then we would expect to observe clustering scores at chance levels (.5).

One-sample *t*-tests revealed that both clustering scores significantly exceeded this level, *t*(51)′*s* > 7.4, *p* < .0001, indicating a tendency to cluster Repeat items with neighboring items from both presentations. However, paired-samples *t*-tests demonstrate that temporal clustering scores were greatest among recalls counted from the serial position of the first presentation, significantly exceeding those of the second presentation, *t*(51) = 14, *p* < .0001. The lag-CRP function (Fig. 2D) confirms this result, showing the strongest probability of transitioning to items that are adjacent to the first presentation. The results indicate that participants were more likely to cluster Repeat item recalls with words presented near the first presentation than the second.

### EEG results

#### Neural evidence of context reinstatement

We next sought to determine whether our EEG recordings captured neural temporal context reinstatement during the recall of Repeat items. Figure 3 presents the ERS values for an item *i* as well as its neighboring items presented at serial positions *i* − 5 to *i* + 5. According to retrieved-context models, temporal context reinstatement should result in decreasing neural similarity as a function of absolute lag in both the forward (positive) and backward (negative) directions (Polyn and Kahana, 2008). To test whether recalling a Repeat item accompanied neural reinstatement of either its first or second occurrence within the list, we examined neural similarity across Lags -5 to 5 for both presentations.

**Figure 3.**
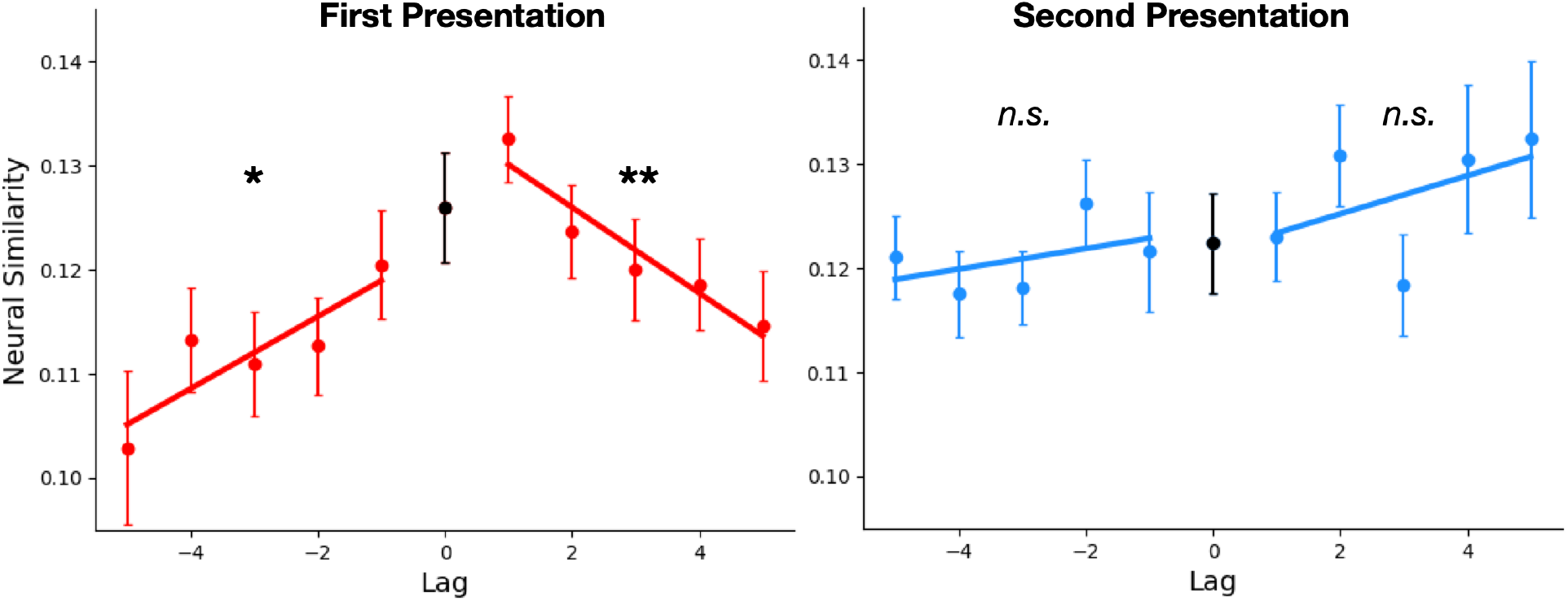
Neural correlates of context reinstatement. ERS for Repeat words and their neighboring study list items. ERS is shown separately for the first (left) and second (right) presentations of Repeat items. Lag refers to the distance in serial position between two items during study. Error bars indicate 95% within-participant confidence intervals (Loftus and Masson, 1994).

Prior studies that measured temporal context reinstatement with neural ERS developed two separate methods for testing the significance of their effects. Lohnas et al. (2023) performed paired-samples *t*-tests comparing ERS values for absolute Lag 1 with those of absolute Lags 3-5 separately across positive and negative lags to verify that neural similarity decreases with serial position distance. Manning et al. (2011) separately computed the across-participants one-sample *t*-statistics for the slopes fit over the positive and negative lags to confirm that similarity values increased over the negative lags and decreased over the positive lags. We submitted our results to both tests, finding that for first presentations (Fig. 3, left), ERS decreased significantly with greater temporal distance over both the negative (*t*(51)′*s* > 1.8, *p*′*s* < .05) and positive lags (*t*(51)′*s* > 2.6, *p*′*s* < .01). However, when we examined ERS for second presentations (Fig. 3, right), we did not observe decreasing neural similarity with temporal distance over either the positive or negative lags, all *t*(51)′*s* < .67, *p*′*s* > .25 (Fig. 3, right). Thus, both statistical tests converge to indicate that we detected evidence of neural context reinstatement for the first presentation, but not the second.

To test whether neural context reinstatement significantly differed between first and second presentations, we constructed a linear mixed-effects model using the lme4 package in R (Bates et al., 2015). The model included ERS as the dependent variable; fixed effects of absolute lag, presentation number (first/second), and their interaction; and random effects of participant nested within presentation number and absolute lag. We created the model using the following notation:

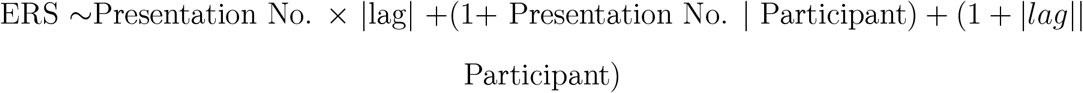

We excluded ERS values for Lag 0 to account for potential item-specific reinstatement effects (Manning et al., 2011; Lohnas et al., 2023). ANOVAs on the model indicated that, although there was no main effect of presentation number, ERS significantly decreased overall with serial position distance, *F* (4, 358) = 4.0, *p* = .04. Importantly, an interaction of presentation number and |lag| revealed larger decreases in ERS as a function of absolute lag for the first presentation than for the second, *F* (4, 933) = 6.5, *p* = .01.

Consistent with our behavioral results, the neural ERS analyses therefore demonstrate stronger evidence for context reinstatement of the initial presentation of a Repeat word.

## Discussion

The present study investigated the relationship between recalling a repeated event and reinstating the neural context from one or both occurrences. Behavioral analyses revealed a tendency to temporally cluster recalls around both presentations of a repeated item. However, contrary to an expected recency advantage, subjects exhibited stronger temporal clustering around the first item presentation. Complementing these behavioral findings, neural activity during the recall of repeated items exhibited graded similarity to activity recorded while encoding items studied near the first presentation, but not the second. Together, these results suggest that people form stronger context associations from an item’s initial presentation than subsequent repetitions, aligning with retrieved-context frameworks of episodic memory.

We innovated a unique combination of behavioral and neural analyses to measure temporal context reinstatement of the repeated words. Specifically, we adapted the lag-CRP function and temporal clustering score, enabling a single recall transition to yield distinct measurements for each presentation of the word. We similarly adapted an EEG signature of temporal context reinstatement to compare neural activity at recall with that recorded near each presentation. Although other measures of temporal context binding exist (Ezzyat and Davachi, 2014; Faber and Gennari, 2017; Broitman and Swallow, 2023), our approach offers the advantage of not explicitly requiring participants to judge temporal relationships between items. This enabled us to observe the influence of repetition on temporal organization in memory during both encoding and retrieval, even when temporal associative memory was not directly relevant to the task (see Lohnas et al., 2023).

Our measures converged to show that when recalling a Repeat word, both behavioral and neural markers of context reinstatement favor its first presentation over the second. Notably, although the encoding-retrieval similarity analysis did not reveal neural context reinstatement for the second presentation, participants temporally clustered their recalls with other items presented near the second presentation significantly above chance levels, albeit at lower levels than the first. This might suggest a divergence between our behavioral and neural measures, however it is more likely that the neural ERS measures lack sufficient sensitivity to detect any subtle reinstatement effects for the second presentation.

Behavioral context reinstatement effects tend to be larger in magnitude than their neural counterparts, with both measures correlating at the individual level (Manning et al., 2011; Lohnas et al., 2023), which could explain why we did not observe neural reinstatement of second presentations. One might further consider whether increased clustering around the first presentation simply reflects better encoding of early list items. If individuals recalled early list items at a higher rate overall, then they would be more likely to successively output these words near a repeated item. In this case, we would expect to observe an asymmetric serial position curve for the once-presented words, with recall favoring primacy items. In contrast with this prediction, we observed a symmetrical SPC function and numerically greater recall for recency items than for primacy items, though this difference was not significant, *t*(51) = 1.8, *p* = .08. However, future studies could further explore this possibility by implementing experimental parameters that modulate primacy and recency effects to investigate their impact on the reinstatement of repeated items.

Studies of proactive interference have demonstrated that associative information from an item’s initial presentation persists longer than that of later occurrences (Briggs, 1954; Underwood, 1957). Such demonstrations run contrary to the law of recency (Kahana et al., 2024) wherein more recent experiences enjoy a mnemonic advantage at all time scales. The mechanism of study-phase retrieval offers a means of reconciling these observations. As argued by Greene (1989) based on a large body of prior research (Melton, 1970; Hintzman, 1976; Thios and D’Agostino, 1976), the second presentation of an item within a list can reinstate its initial presentation. Greene suggested that study-phase reinstatement may underlie the beneficial effects of spacing on recall of repeated items, a finding that also runs counter to a recency-based account (because moving the first presentation of a repeated item farther into the past results in better rather than worse memory). As a test of study-phase reinstatement, Lohnas and Kahana (2014) found that subjects were more likely to successively recall words presented immediately after each instance of a repeated item as compared with equally spaced non-repeated items. They interpreted this result in terms of retrieved context theories of memory: study-phase retrieval leads to greater similarity among the contexts associated with repeated items, causing retrieval of neighbors from one presentation to evoke the neighbors of the other. Research using fMRI and intracranial EEG has further revealed that repeating an item within a list reproduces neural patterns observed at the item’s initial occurrence in a manner consistent with study-phase context reinstatement (Xue et al., 2010; Halpern et al., 2023). The current study extends this research by demonstrating that individuals cluster the retrieval of repeated items with other words presented near its initial presentation and that recalling a repeated item accompanies the reinstatement of neural patterns observed near the first presentation but not the second, possibly indicating that study-phase reinstatement strengthens preexisting item-context associations. Together, these findings support CMR’s assumption that repetition effects arise from a combination of contextual variability across multiple presentations and study-phase reinstatement of prior occurrences during repeat presentations.

Although CMR accounts for our results, alternative explanations also merit consideration. One such theory posits that detecting a repeated item suppresses encoding in a process known as *repetition-suppression* (see Auksztulewicz and Friston, 2016 for a review). Repetition-suppression could weaken the activation of an item and its corresponding association with the context of the second presentation. If an item is successfully encoded upon its first presentation, then individuals might predominantly rely on representations from that initial occurrence during retrieval. Our data support this notion: participants in our study successfully recalled approximately 56% of Single items (Fig. 1A), suggesting a 56% chance of encoding success per presentation. If this were the case, and participants successfully encoded 56% of Repeat items during the first presentation, then they might retain an additional 25% from the remaining 44% of Repeat items upon their second presentation. This would produce a recall rate of 81% (we observed an average rate of 79%). Notably, 69% of the Repeat items recalled would have been encoded during their initial occurrence, retaining context associations from that first presentation. Thus, repetition suppression may not only explain the difference in recall rates between Single and Repeat items, but also our observation of increased reinstatement of the first presentation. However, further investigation is needed to test this theory, particularly through experimental conditions that reduce the likelihood of successful encoding from a single presentation (e.g., by speeding the presentation rate) to assess whether this leads to greater reinstatement of later presentations.

### Study limitations

Several aspects of the current experiment’s design may have constrained our ability to distinguish temporal context reinstatement between the first and second presentations of Repeat words. Specifically, presentation position (first vs. second) was confounded with serial position, as second presentations occurred later in the study list than first presentations. This could have affected the neural ERS analyses for the second presentation of repeat items, as there were fewer opportunities to examine ERS for lags 1 to 5 for these later occurrences in the final positions of the study list, or for lags -5 to -1 for first presentations that occurred at the start of the list. Additionally, since we examined neural activity from up to five serial positions away but some repetitions were separated by only three lags, the presentations sometimes overlapped in the ERS analysis, leading to their exclusion from our results. Future studies might mitigate these issues by extending list lengths and ensuring all repetitions are spaced beyond the maximum lag analyzed for ERS. Further limiting our analyses, we did not present a single item more than twice within a list in the current study. As a result, we cannot definitively resolve whether temporal context reinstatement favors the initial presentation of an item or the one preceding its last occurrence. Future work could address this by presenting some items more than twice within a list or repeating items across lists (e.g., Zaromb et al., 2006).

### Conclusions

Our daily lives are filled with recurrent experiences. Although such events occur within distinct temporal contexts, forming shared representations facilitates learning and generalization. Here we found evidence that, in contrast with recency-based accounts of context reinstatement, participants neurally reinstate temporal context information from the first instance of a repeated event, but not the second, during recall. Together, these results indicate that people form stronger temporal context associations with the initial occurrence of an event than with later repetitions.

